# Relative Concentration of Brain Iron (rcFe) Derived from Standard Functional MRI

**DOI:** 10.1101/579763

**Authors:** Stan J. Colcombe, Michael P. Milham, Anna MacKay-Brandt, Alex Franco, F. Xavier Castellanos, R. Cameron Craddock, Jessica Cloud

## Abstract

Brain iron plays key roles in catecholaminergic neurotransmitter synthesis and early life brain development. It is also central to cellular energetics and neurotransmitter metabolism throughout the lifespan. Disturbances in brain iron have been implicated in a growing number of psychiatric and late-life neurodegenerative disorders. Additionally, brain iron accumulations are thought to play a deleterious role in neuroinflammatory processes in later life. Despite its importance, the role of brain iron in development, aging, and psychiatric disorders remains comparatively understudied. This is partly due to technical challenges inherent in implementation and analysis of formal iron imaging protocols and practical constraints on scan session durations. Here, we introduce a method to estimate relative brain iron concentrations that is 1) computationally simple, 2) shows excellent correspondence with formal iron imaging in-vivo, 3) replicates clinically-relevant findings from formal iron imaging, 4) yields novel insights into brain iron and cognition across the lifespan, and 5) leverages a widely available and frequently shared brain imaging modality: functional MRI. The computationally simple nature of the measure, coupled with the availability of fMRI datasets across the lifespan and disorders, has the potential to transform our understanding of the complex and critical relationship between iron and brain health.

## Introduction

Iron metabolism and accumulation are undisputedly essential to healthy brain function, but their roles in brain health are complex when considered across the lifespan. In prenatal life, iron serves to guide neuronal development, subserves myelination^1^ and dopamine receptor development^2^, and is essential for synthesis of catecholaminergic neurotransmitters. This is particularly evident throughout childhood^3^ and into later life in areas such as the striatum, midbrain, and brainstem areas such as the substantia nigra, ventral tegmental area and locus coeruleus^4,5^. As in most cells in the mammalian body, iron also plays a central role in brain cellular energetics^6^, as well as myelin maintenance throughout the lifespan^1,7^. Iron is poorly absorbed into the body, tightly regulated once incorporated, and crosses the blood-brain barrier via active transport^8,9^. Unlike most other metals, active mechanisms to eliminate iron from the body do not exist^10^. Indeed, in areas where iron utilization is high, such as the striatum and other catecholamine-rich areas, iron tends to gradually accumulate throughout the lifespan^11,12^. Unfortunately, accumulation of iron results in cellular oxidative stress^13^, suggesting a potential deleterious role of brain iron accumulation in otherwise non-pathological aging.

It is also, perhaps, not surprising that dysregulation of this key element is increasingly associated with a number of psychiatric disorders. Some, such as restless leg syndrome^14^, and Attention-Deficit Hyperactivity Disorder^14,15^ (ADHD) are associated with reduced brain iron levels, with evidence that iron supplementation can provide some symptom relief^16–18^. Other disorders such as Parkinson’s disease^19–21^, and Alzheimer’s dementia^20,22,23^ are associated with excessive regional brain iron, leading to hypotheses about ferroptosis (iron-mediated cell death)^4,24–26^, and approaches to reduce iron levels as a target for intervention^20,24,27^ Given these observations, iron appears to be a useful index to track normative maturation from early to late-life, while also serving as a reliable indicator of pathological disruption from that normative trajectory.

Unfortunately, despite the clear importance of brain iron, both as a boon for brain development and metabolism and as a hazard through later life oxidative stress, iron has been comparatively understudied with neuroimaging, particularly relative to modalities such as functional Magnetic Resonance Imaging (fMRI). This can likely be ascribed to the requirements for specialized imaging sequences in MRI studies, which tend to be deprioritized relative to functional, diffusion, and morphometric imaging protocols, as well as specialized knowledge required to analyze standard quantitative iron imaging protocols. These observations highlight the need for a computationally tractable method to quantify brain iron that can be generated without compromising or overburdening ongoing data collection efforts.

Here, we report findings from targeted investigations examining whether standard fMRI images can be leveraged to estimate relative brain iron concentrations. We conclude that our approach, termed relative concentration of brain iron (rcFe), has substantial potential to facilitate the study of brain iron in development, aging, neurologic and psychiatric populations. We argue from first principles that fMRI sequences should be sensitive to brain iron concentrations, and proceed to report a series of findings supporting this claim and its intriguing implications. However, in the interest of transparency, we note that our results emerged serendipitously from explorations of possible confounds in the quality of fMRI skull stripping algorithms as a function of participant age. After resolving the skull stripping issue, the lead investigator had a large sample (n=341) of skull-stripped and coregistered 3D mean fMRI images. Out of curiosity, he regressed age against mean fMRI value at each voxel. The pattern of results was striking, resembling well-established patterns of iron accumulation in the striatum across the lifespan. What follows is a report of our attempts to refine these initial observations into a workable method and explore its utility.

Our rcFe approach builds upon the well-established relationship between the paramagnetic properties of unbound iron and its impact on T2* image susceptibility. This general property of susceptibility is fundamental to imaging protocols designed to measure tissue iron levels ^5,28,29^, as well as fMRI^30,31^. A typical approach to quantitative iron imaging involves the collection of structural T2- or T2*-weighted images at varying delays in echo time (TE). Higher regional brain iron levels increase regional susceptibility, and therefore the rate at which the signal decays over time. It is then possible to fit a decay function to quantify iron content at each voxel ^32–34^. Similarly, fMRI sequences capitalize on the fact that deoxygenated blood is paramagnetic and increases the rate of signal decay, much like unbound iron, while oxygenated blood is moderately diamagnetic and has the effect of slowing the rate of signal decay^35^. However, standard fMRI sequences use a fixed TE that optimally distinguishes oxygenated vs. deoxygenated blood levels (i.e., blood oxygen level dependent, or BOLD signal). fMRI sequences only collect a single TE, and therefore cannot be used to estimate a decay function at each voxel. However, they are, by design, sensitive to variations in magnetic susceptibility such as those caused by variations in brain iron. Although it is not possible to estimate absolute iron content from a susceptibility image with a single TE, it may be possible to provide an estimate of the relative iron content at each voxel, particularly for regions that are high in iron content such as the striatum. As an example, Figure 1, top, demonstrates how the signal in a susceptibility weighted image might decay given different levels of iron when sampled at 7 different echo times; these values would be fit to an exponential decay function; the reciprocal of this fit is linearly related to iron content (e.g., 1/T2=R2; 1/T2*=R2*). Figure 1, bottom, illustrates how those same decay curves might be sampled at a single TE (e.g., 2E), and inverted to create an estimate of the relative concentration of brain iron, which we term rcFe.

**Figure 1.**
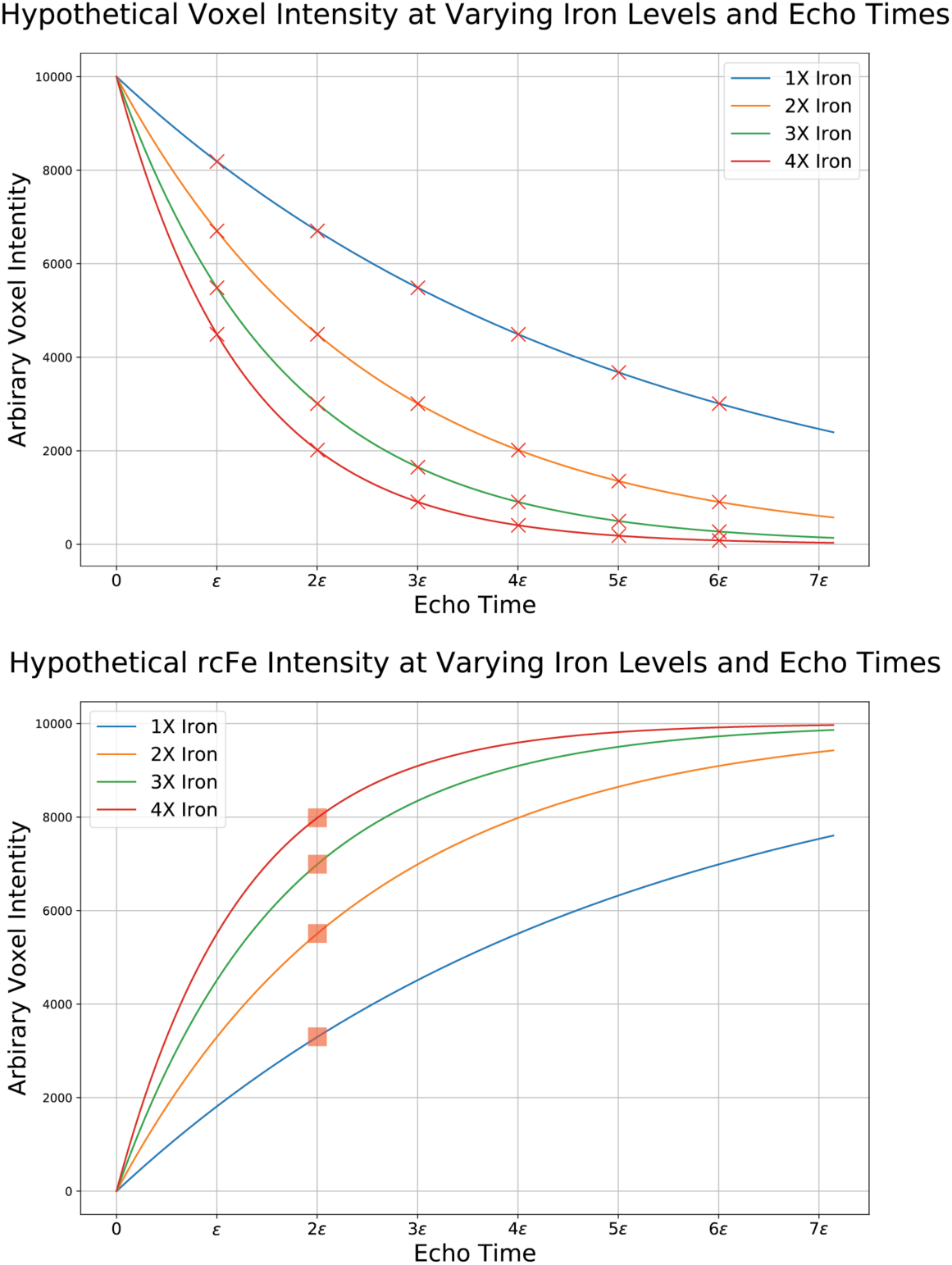
Top panel: illustration of expected signal decay in a susceptibility-weighted image at varying levels of iron content. Bottom panel: the same curves as in the top panel, but inverted to show how the expected signal decay at varying iron concentrations might be represented in a single-echo susceptibility image, an approach we have termed relative concentration of brain iron (rcFe).

In this paper, we first describe our initial finding of age-related variability in mean fMRI signal. We then report on a series of studies carried out after observing the striking consistency of our initial findings with extant quantitative MRI and post-mortem studies of brain iron concentrations. Specifically, we report on our efforts to: 1) explore variability in our rcFe measure by evaluating a large lifespan sample through independent components analysis (ICA), we then 2) validate the sensitivity of rcFe to iron in an aqueous phantom containing iron levels selected to approximate known iron concentrations within the human brain, 3) examine the correspondence between rcFe measures and standard approaches to iron imaging, 4) replicate established findings related to subcortical iron concentrations in a clinical population, as well as reveal a novel finding in the relationship between brain iron and ADHD symptomatology in typically developing children, and 5) provide novel insights into the age-varying relationship between relative subcortical iron concentrations and IQ across the lifespan.

## Results

### Initial observations

The initial findings demonstrating the impact of age on the mean fMRI signal are shown in Figure 2, left panel. The peak clusters, thresholded at Z>18, clearly highlight the anatomical structure of the putamen. The overall relationship between age and mean fMRI signal in the displayed mask plotted in Figure 2, right panel, illustrate a striking relationship between age and mean fMRI signal (r(339)=0.81, p<0.001). This general observation is striking not only in the strength of the relationship, but also in that the peak effect centers on an area that is amongst the most intensely concentrated with iron in both MRI and post mortem studies^28,29,36,37^, and the consistency with the predicted impact of iron on a susceptibility weighted image such as fMRI.

**Figure 2.**
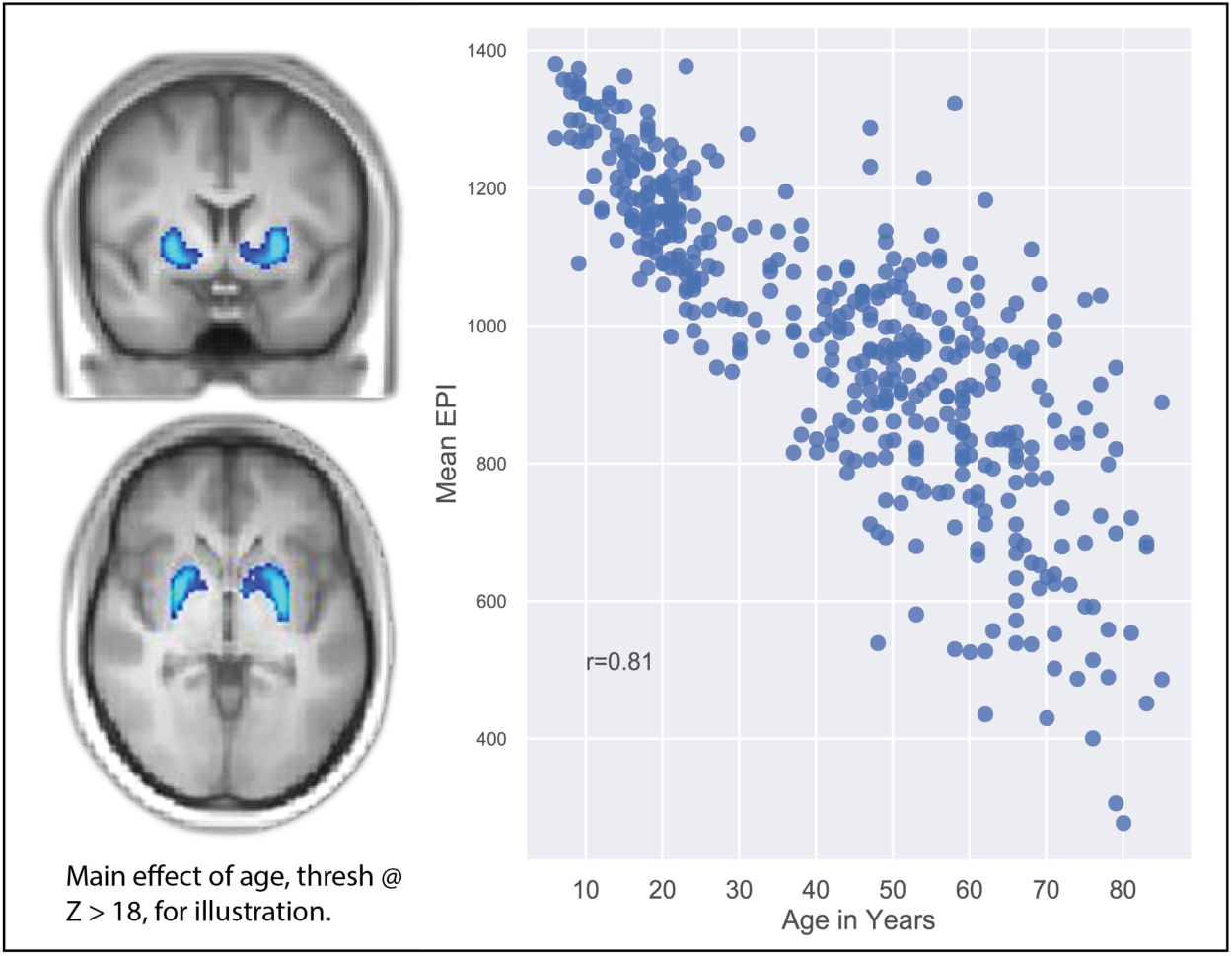
Initial observations. The results from a serendipitous assessment of variation in mean fMRI signal across the lifespan. Peak variation was apparent in the iron rich putamen (left). The mean within-mask values are plotted in the right panel.

### Independent components analysis in larger NKI-Rockland Sample Initiative

Our initial observation related to regional age-wise variation in mean fMRI signal was provocative and consistent with predictions that might be made regarding the accumulation of brain iron across the lifespan. A further exploratory ICA investigation of the rcFe images within a larger NKI-RSI cross-sectional sample yielded additional insights into the patterns of variation across the lifespan. Inspection of the ICA outcomes revealed three distinct spatial components encompassing three iron-rich subcortical structures. These components are shown in Figure 3, top panel, and include aspects of the putamen and caudate (red), globus pallidus (blue), and thalamus (green), overlain on the MNI152 T1 template. Figure 3, lower panel shows the mean rcFe values within those three ROIs plotted against age. rcFe values were significantly correlated with age in all three ROIs, although the putamen (r(1352)=0.72) and globus pallidus (r(1352)=0.62) correlations were notably stronger than that for the thalamus (r(1352)=0.15); all ps <0.0001. Again, these findings are generally consistent with previous MRI and post-mortem studies of striatal brain iron accumulations.

**Figure 3.**
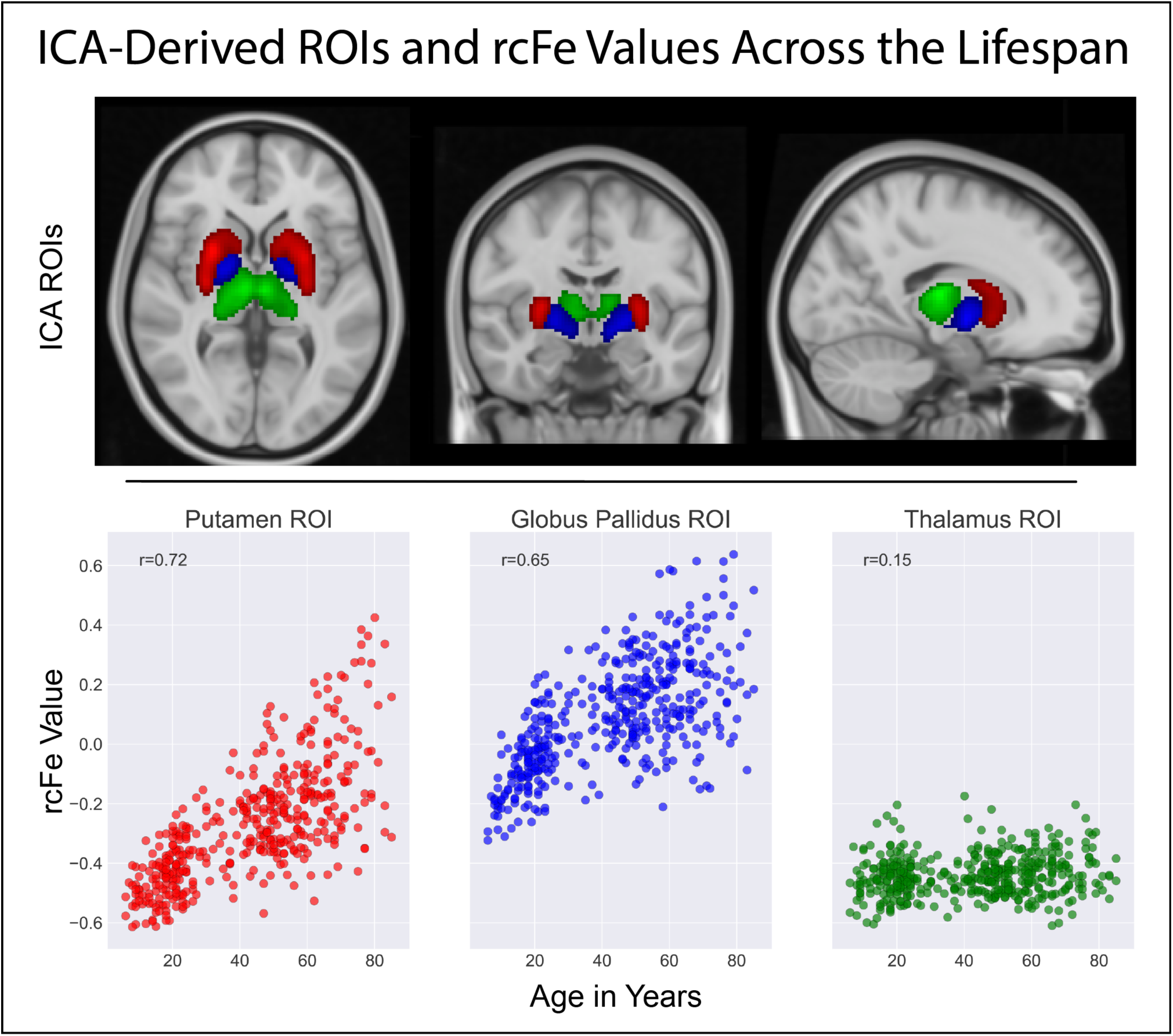
Top panel: Independent components analysis of rcFe values revealed three subcortical components, putamen and caudate (RED), globus pallidus (BLUE), and thalamus (GREEN). Bottom panel: Mean rcFe values for each of the three ROIs plotted against age for each of the 1354 participants.

### Aqueous iron phantom

To examine the sensitivity of rcFe images to known iron concentrations, we examined rcFe values from a phantom with varying aqueous iron concentrations, scanned under a well-established fMRI sequence. Figure 4, inset, shows a photograph of the aqueous phantom. Figure 4 shows a stripplot of all 216 rcFe voxel values for each iron concentration level, taken from 6×6×6 voxel regions of interest placed near the spatial center of the vials containing each iron concentration in the phantom. Voxel rcFe values for each iron concentration within the ROIs are non-overlapping, with reduced variance at higher iron concentrations. Overall, these results bear a striking resemblance to the expected pattern of results from a hypothetical single TE susceptibility weighted image, and appear to confirm the sensitivity of rcFe to biologically plausible iron concentrations.

**Figure 4.**
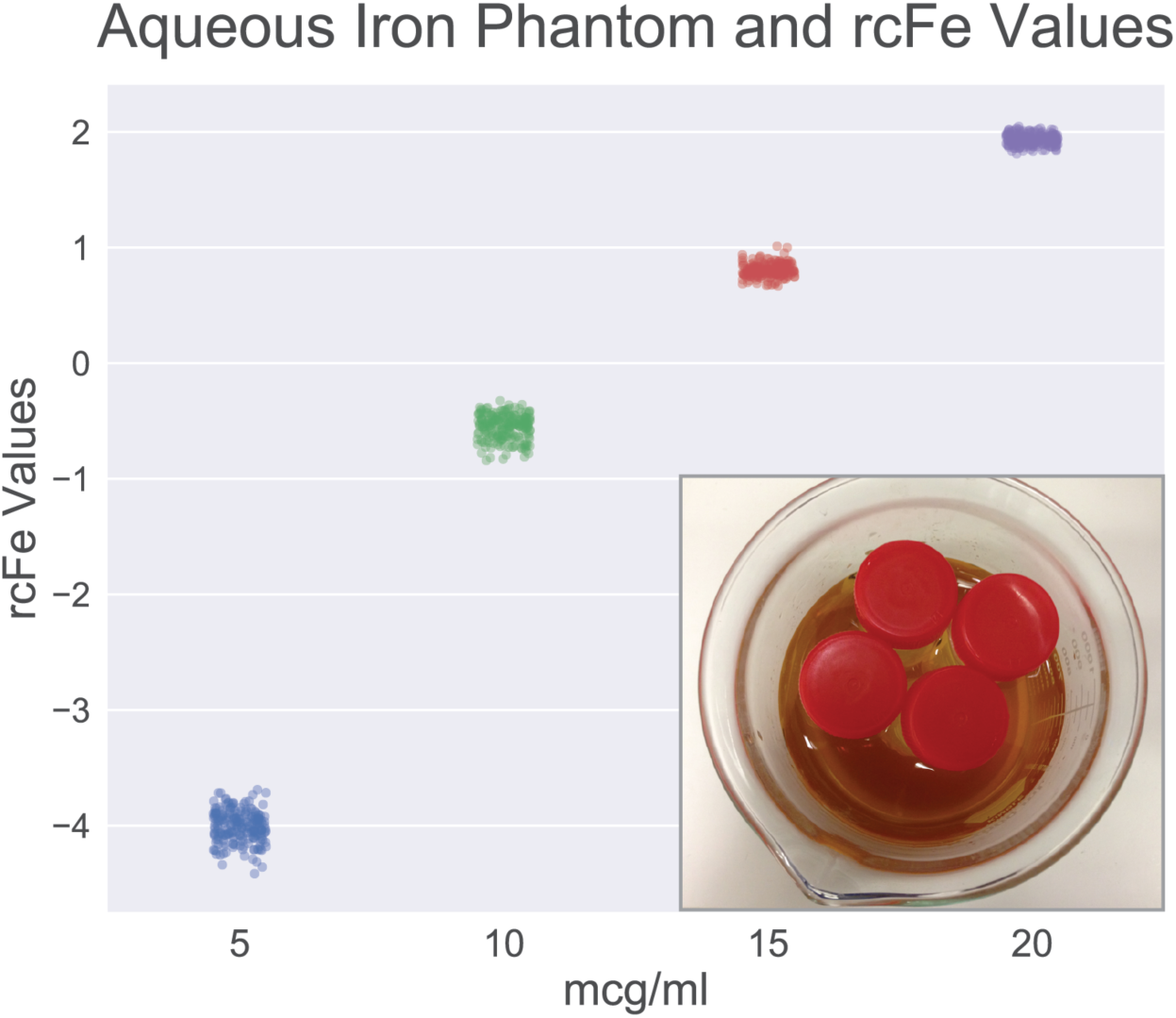
Inset: Aqueous iron phantom used to assess the sensitivity of rcFe to known iron concentrations selected to be biologically plausible. Stripplot showing the rcFe values for all 216 voxels extracted from the center of each iron phantom vial.

### In-Vivo comparison between rcFe and formal iron methods

Figure 5 illustrates the relationship between rcFe and iron levels estimated from three different iron imaging protocols in 10 subcortical regions of interest taken from the Harvard-Oxford subcortical atlas. The top panel plots the rcFe values against the normalized iron values for each ROI for 5 participants. Each colored line represents a best fit line for each participant. The overall correlations between ROI values for rcFe and the formal iron imaging protocols within subcortical structures was robust, with *r*(48)=0.92, r(48)=0.93, r(48)=0.84 for the Siemens clinical, 10-echo R2, and 6-echo R2* sequences, respectively; all p values < 0.0001. The lower panel shows a 25% subsampling of within-brain voxel-values for rcFe plotted against values derived from each of the three iron imaging sequences for a single representative participant. These correlations were substantially lower than those constrained to subcortical regions of interest, with r=0.26, r=0.29, and r=0.27, for the Siemens clinical, 10-echo R2, and 6-echo R2* sequences, respectively; all ps < 0.0001. Overall, this pattern of findings suggests that there is good correspondence between rcFe and some more traditional iron imaging approaches, at least within well-defined subcortical areas. These findings, in combination with those from the aqueous iron phantom, suggest that rcFe has reasonable sensitivity to relative iron concentrations.

**Figure 5.**
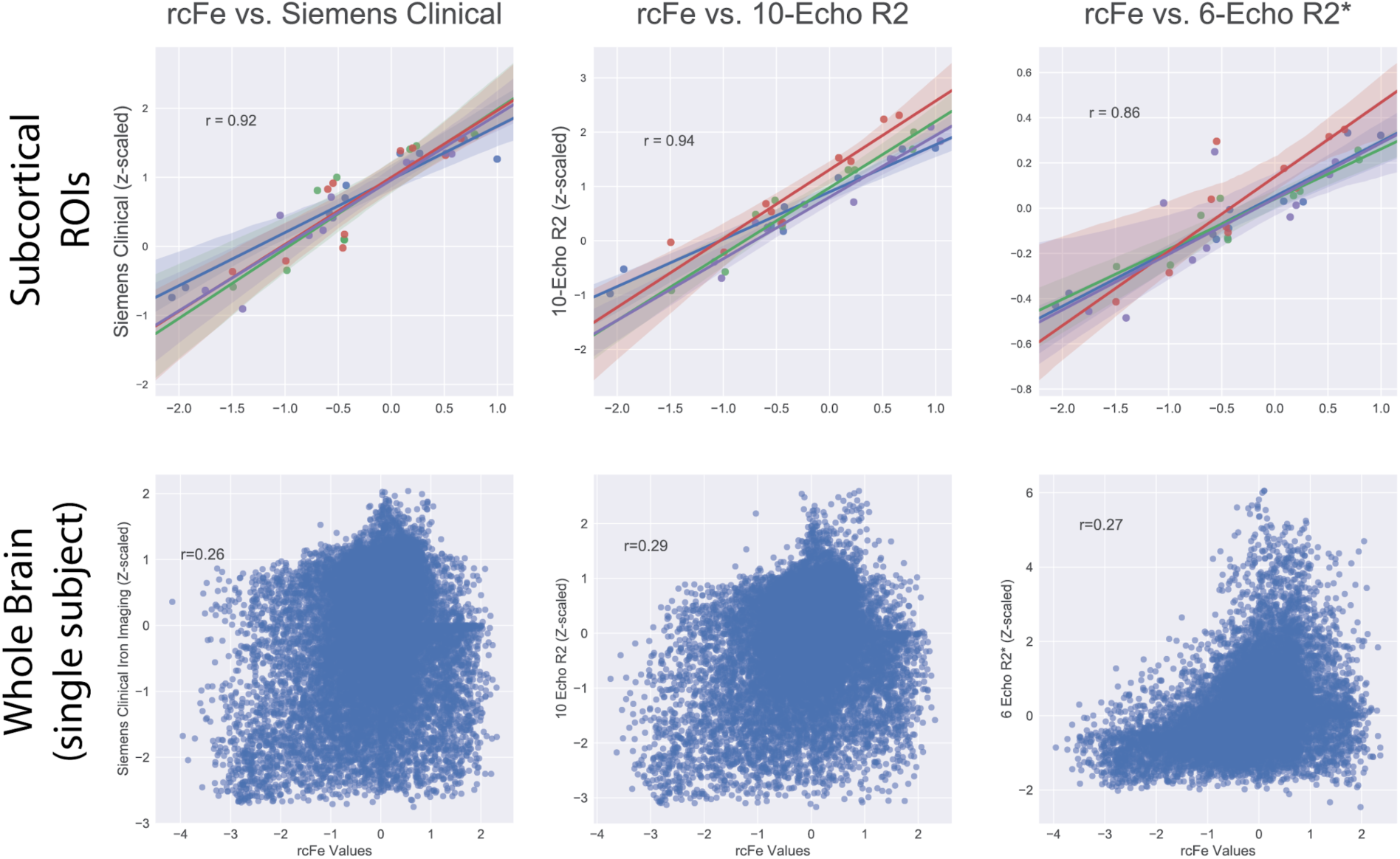
In-vivo comparison between rcFe and three standard iron imaging approaches. Top panel: Mean rcFe and iron values for each of 10 subcortical ROIs taken from the Harvard Oxford Subcortical Atlas. Bottom panel: voxelwise plots of rcFe and iron estimates for a single subject.

### Examination of rcFe clinical relevance within the ADHD200 Sample

#### Conceptual replication of NKI-RSI ICA

Having established the sensitivity of rcFe to iron content in the aqueous phantom and good correspondence between rcFe and more standard assessments of brain iron, we next examined the utility of rcFe in a clinical population. Specifically, we examined rcFe in an openly available sample of ADHD and typically developing children (TDCs), given the well-established reductions in brain iron associated with ADHD. As an initial examination, we conducted a conceptual replication of the ICA analysis (reported above) from the NKI-RSI dataset. Specifically, rcFe images were computed from the ADHD200 dataset, preprocessed, and submitted to exploratory ICA. Visual inspection of the resulting component maps readily identified three subcortical components showing a striking similarity to those identified in the NKI-RSI dataset. Figure 6 shows these components overlain on the MNI152 T1 template; again these include aspects of the putamen and caudate (red), globus pallidus (blue), and thalamus (green).

**Figure 6.**
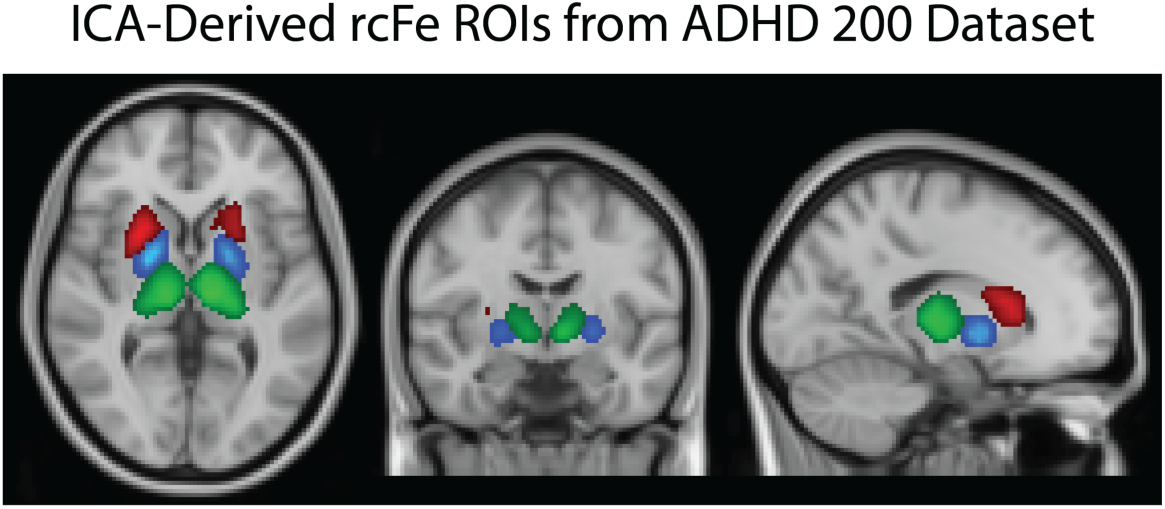
Conceptual replication of NKI-RSI ICA findings. Similar subcortical components are evident in the ADHD200 sample to those identified in the NKI-RSI.

#### Group differences in subcortical iron

Perhaps not surprisingly, initial comparisons of ROI values revealed a significant effect of age in the putamen (F[1,616]=5.67, p<0.018), globus pallidus (F[1,616]=,7.91, p.<0.005), and thalamus (F[1,616]=8.16, p.<0.004). However, there were also substantial site effects for putamen (F[1,616]=68.75, p<0.001), globus pallidus (F[1,616]=99.04, p.<0.005), and thalamus (F[1,616]=8.16, p.<0.004), suggesting that z-scaling rcFe values within-volume is not sufficient to completely remove sitewise differences in hardware or scanning protocols. Importantly, however, rcFe was significantly lower for ADHD participants, compared to controls in the putamen (t[616]=6.97, p.<0.0001), globus pallidus (t[616]=5.25, p.<0.0001), and thalamus (t[616]=4.99, p.<0.0001). See Figure 7. Moreover, accounting for the effects of age and site in the contrast between ADHD and TDCs resulted in a more robust effect of diagnosis for putamen (t[614]=8.61, p.<0.0001), globus pallidus (t[614]=7.07, p.<0.0001), and thalamus (t[614]=6.82, p.<0.0001), suggesting that while sitewise differences in protocol may contribute noise to datasets pooled across disparate sites and protocols, the overall effect of rcFe-estimated iron content in ADHD vs. TDCs was quite robust and consistent with expectations regarding typically iron-deficient ADHD participants.

**Figure 7.**
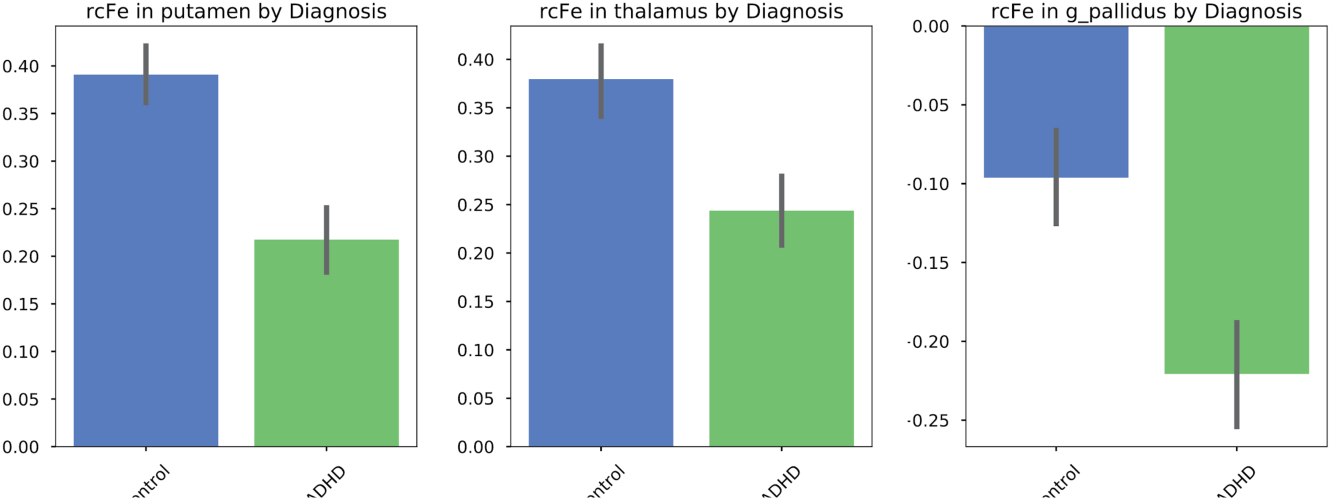
Bar graph showing rcFe values for children with ADHD and TDCs. These findings are consistent with previous observations of depleted brain iron levels in ADHD.

### ADHD symptom severity in ADHD and TDC

#### ADHD symptom severity and subcortical rcFe

The overall pattern of group differences in diagnostic status was consistent with the idea that participants with ADHD had lower subcortical brain iron than TDCs. However, iron deficiency has been proposed as a causal mechanism for ADHD symptomatology^38^, symptom severity appears to be modifiable by iron supplementation, and symptom severity is correlated with serum iron levels in blood^38^. This suggests that estimated brain iron levels may predict ADHD symptom severity along a continuum.

To investigate this possibility, we restricted analyses to participants from the ADHD200 sample who had valid ADHD symptom scores on the ADHD Rating Scale IV (ADHD-RS)^39^ provided with the ADHD200 phenotypic information; this resulted in a sample of 374 participants (155 ADHD, 219 TDC). No significant difference in age existed (11.7 and 12.01 years for ADHD and TDCs, ns.). There was, as expected, a significant difference in adhd symptom severity score as a function of diagnostic status. Participants with ADHD scored significantly higher on symptom severity than TDC participants (57.1 vs. 37.5, respectively; t[372]=11.37, p<0.001).

We entered diagnostic status (ADHD vs TDC) and mean rcFe values into separate regression models predicting ADHD symptoms for each of the 3 ROIs^40^. However, given the impact of site and age on mean rcFe value seen above, we first residualized the rcFe values for each ROI by regressing out site and age. Regression analyses revealed that, collapsing across diagnostic status, rcFe was significantly related to ADHD symptomatology for the putamen (F[3,374]=44.01, p.<0.001), globus pallidus (F[3,374]=34.12, p.<0.001), and thalamus (F[3,374]=33.51, p<0.001). Interestingly, however, a significant interaction between diagnostic status and rcFe value predicting ADHD symptom severity for putamen (F[3,374]=24.95, p.< 0.001), globus pallidus (F[3,374]=6.69, p.<0.01), and thalamus (F[3,374]=4.15, p.<0.042). Figure 8 shows the rcFe values plotted against ADHD symptom score, split by diagnostic status. In all three ROIs, it is apparent that the symptom severity overall is negatively related to rcFe intensity, with the suggestion that with reduced rcFe estimated iron levels ADHD symptom severity increases. This is generally consistent with the hypothesis that depletion of subcortical iron is related to ADHD symptomatology. It is also apparent from the plots in Figure 7 that symptom severity is related only to variation in rcFe values in the TDC sample.

**Figure 8.**
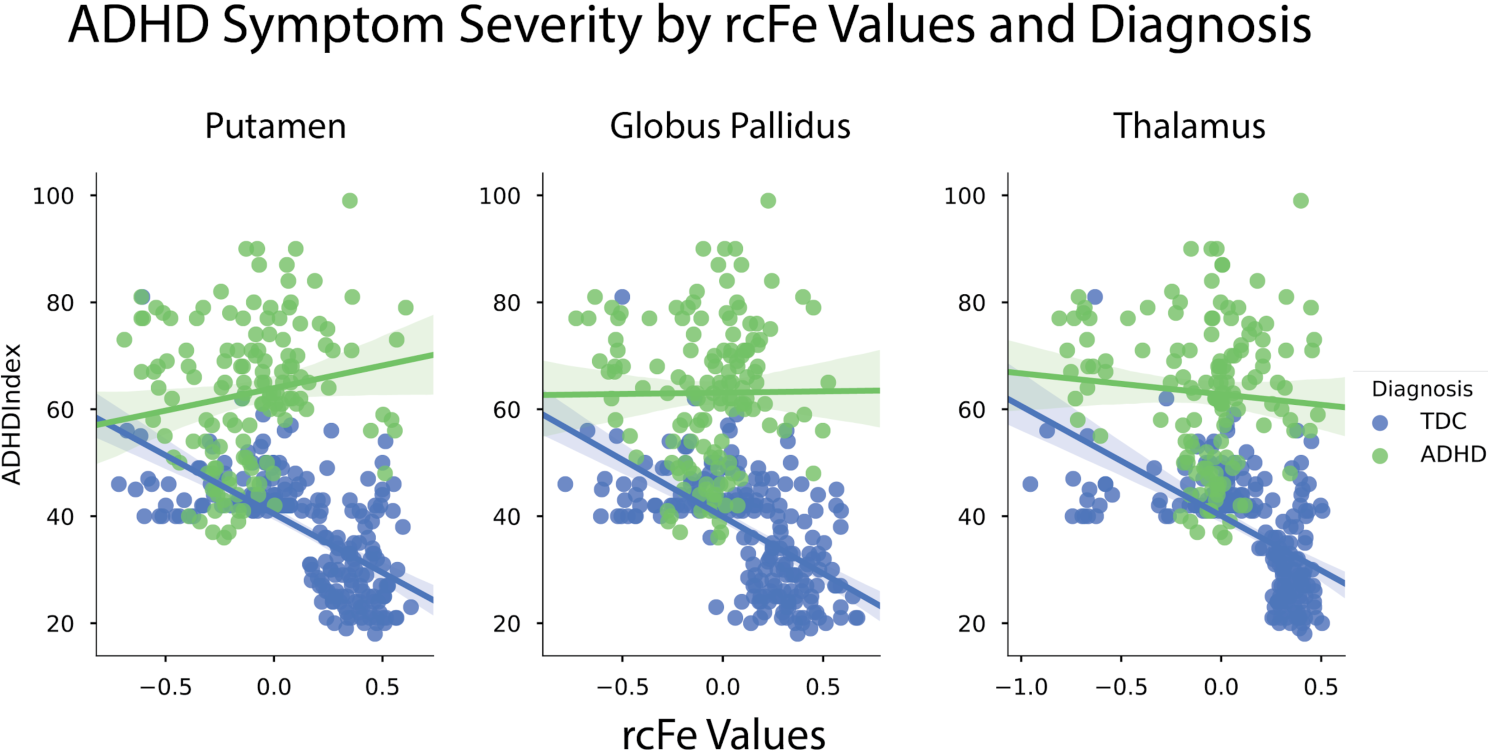
rcFe values for each of the 3 subcortical ROIs, corrected for age and site, plotted against ADHD symptom severity separately for children with ADHD (Green) and TDCs (Blue).

### rcFe, age, and IQ in the larger NKI-RSI

Within a developmental sample, it appears that reduced brain iron is detrimental to brain health, while later in life accumulation of brain iron appears to be associated with oxidative stress and a host of neurodegenerative disorders. We returned to the NKI-RSI to address the potential implications of brain iron and its effects on brain health from a lifespan perspective. We again leveraged the 3 subcortical ROIs identified in the NKI-RSI, and conceptually replicated in the ADHD200 sample. Specifically, for each of the ROIs we examined the relationship between putative regional brain iron as assessed by rcFe and full-scale IQ as a function of age, using a varying coefficient model framework^41^. We modeled the relationship between full-scale IQ and rcFe in each of the three subcortical ROIs, examining the varying effect of age on that relationship, assuming a gaussian smoother function and identity linkage. Figure 9, top panel, shows the age-varying relationship between rcFe estimates of brain iron and IQ across the lifespan. In each of the three ROIs, higher brain iron is associated with better performance on the IQ test in childhood, while in middle and later life rcFe estimates of brain iron levels become irrelevant or detrimental to IQ performance. The age by iron smoother terms predicting IQ were statistically significant in the putamen (F[9.48, 1304]=54.4, p<0.0001), globus pallidus (F[8.5, 1304]=26.7, p<0.0001) and thalamus (F[9.7, 1304]=51.9, p<0.0001).

**Figure 9.**
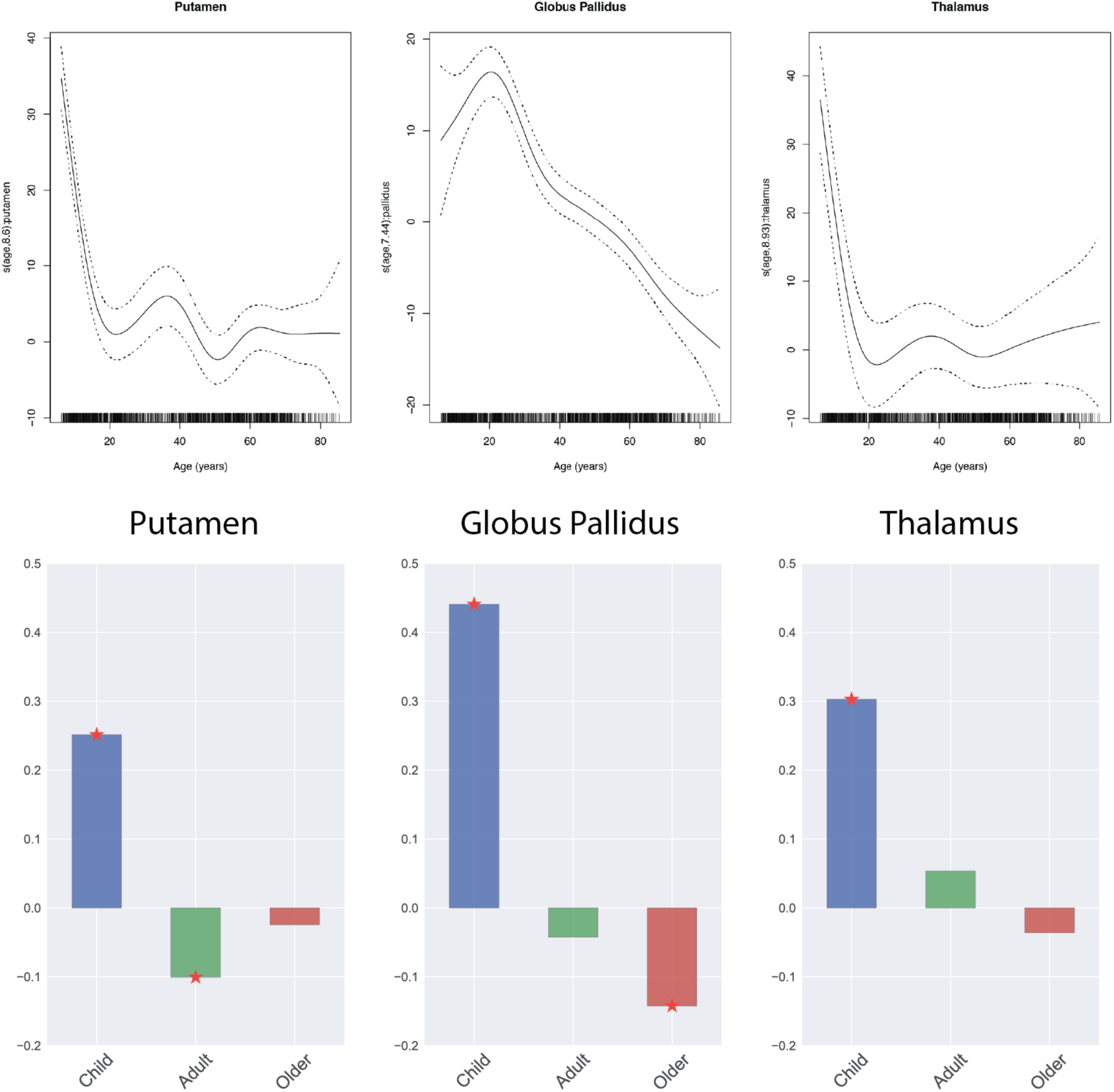
Top panel: Relationship between rcFe and IQ, as a function of age. Bottom panel: A simplified representation of the varying coefficient model above, showing correlations between rcFe and IQ for child, adult and older participants. In both cases, it is apparent that rcFe and IQ are positively correlated in childhood, but either irrelevant or detrimental in adulthood. This pattern is consistent with the opposing effects of brain iron and brain health in development, compared to aging.

Figure 9, lower panel shows a simplified representation of these effects by presenting the correlation between rcFe ROI values in each of three age bands: Child (6-17), Adult (18-45), and Older (46-85). In all three ROIs, the child age band demonstrated a significant positive correlation with rcFe brain iron estimates (ps <0.01). Notably, however, that pattern was significantly reversed in the putamen for adults, and globus pallidus ROI older adults (ps. < 0.01).

## Discussion

Despite the clear importance of brain iron in development, aging, and psychiatric disorders, it remains relatively understudied. Here, we presented evidence in support of rcFe, a computationally tractable method to assess relative brain iron concentrations that avoids overburdening or compromising ongoing efforts. The current study resulted from an initial serendipitous observation that resembled well-established patterns of brain iron accumulation across the lifespan. We argued that the idea that mean fMRI images are sensitive to brain iron content has reasonable construct validity given the nature of fMRI sequences and of unbound iron in a magnetic field. We suggested a simple method to compute a marker that would correct for variability in site-wise or individual scaling of an fMRI image, and intuitively convey potential relative iron content. We then validated this approach with an aqueous phantom of water vials titrated to biologically plausible levels of iron, as well as in-vivo subcortical regions compared against three common iron imaging approaches. We also demonstrated the emergence of three subcortical regions through independent components analysis in two independent samples (NKI-RSI and ADHD200). We then leveraged these ROIs to evaluate the relationship between rcFe-based estimates of brain iron in both samples. We conceptually replicated previous findings demonstrating reduced brain iron content in participants with ADHD compared to TDCs using rcFe. We demonstrated predictive validity of the rcFe estimate to classify the sample by diagnosis (ADHD or TDC).. In addition, we found that rcFe was significantly related to ADHD symptomatology in the TDC sample however, this association was not observed in the ADHD sample. Finally, we presented a novel finding demonstrating that the relationship between rcFe and IQ is critically dependent on age, with higher levels of estimated brain iron being beneficial early in life, but either irrelevant or detrimental later in life. This pattern of results is highly consistent with predictions from models of iron and its impact on brain in development vs later life.

The clinical and lifespan findings reported here demonstrate the potential utility of rcFe as an indicator of relative brain iron concentrations. The public availability of large-scale fMRI datasets such as the NKI-RSI, Healthy Brain Network, Human Connectome Project, UK Biobank, and ADNI, underscores the potential for broad-scale reanalysis of existing fMRI data to advance the study of brain iron across multiple dimensions. In particular, rcFe could be a powerful tool to identify early and midlife markers for later life pathological processes that may elucidate environmental and genetic contributions to both normal aging and age-related disease. It should also be noted that our ADHD results, derived from the ADHD200 dataset, demonstrate the power of aggregating rcFe datasets across disparate sites, hardware configurations, and imaging protocols. As such, rcFe also holds promise in the application to a growing number of ad-hoc aggregations of fMRI datasets acquired from clinical populations, including disorders where iron has been implicated in the etiology (e.g. autism [ABIDE], schizophrenia [COBRE]).

While the initial findings with rcFe and its potential to open new avenues of inquiry to brain iron are promising, there are limitations to both the method and the current study. The most obvious limitation of rcFe is that, unlike quantitative approaches, rcFe can at best provide only a relative estimate of brain iron. In the current study, we restricted our analyses to subcortical structures, which by nature have high iron concentrations. The utility of the rcFe method to cortical areas, or even other subcortical areas, remains unexplored. Our approach to calibrating the rcFe by a linear scaling factor worked reasonably well as a first-approximation, but is almost certainly suboptimal. This was readily apparent in the robust site-wise rcFe effects in the ADHD200 sample. Future work evaluating other approaches that better match the predicted signal decay at a fixed TE (e.g.inverse log) are likely to prove fruitful. Unclear what range of parameters might be appropriate (e.g. variation in TE), or specifically how they might affect the contrast seen here. This is highlighted by the robust main effect of site on rcFe intensity within the ADHD200 sample. Future investigations into this question are likely to be highly fruitful. Similarly, despite the successful aggregation of rcFe across the ADHD200 datasets, it remains unclear what limitations exist in fMRI sequence parameters appropriate for rcFe generation. Future work addressing sequence parameter variation (e.g. TE, bandwidth), combining rcFe, quantitative iron imaging, and phantoms of known iron concentration could provide useful information about the impact of sequence parameters, and perhaps even generate an effective post-hoc calibration algorithm for a range of fMRI sequence permutations.

## Materials and Methods

### Initial observations

341 MRI datasets and basic phenotypic information (age, sex), were drawn from the publicly available NKI-RSI sample^42^, a large scale community ascertained dataset that includes extensive imaging and phenotypic assessments. Participant ages ranged from 6-85 (mean = 44.1yoa, 196 females), as a methodological examination of variation in fMRI image skull stripping efficacy as a function of age. Each dataset consisted of the NKI-RSI 1400ms multiband 4 (MB4) resting state fMRI sequence (TR=1400ms; TE=30ms, alpha=80, 64 slices; FOV=224; acquisition voxel size = 2.0mm iso, multiband factor = 4; duration = 10min) and a high resolution T1 structural image (MPRAGE, TR=2500ms; TE=3.5ms; TI=1200ms; voxel size: 1.0mm iso; alpha=8; 174 slices, FOV=256mm; Grappa=2). Images were subjected to the following procedures: fMRI images were motion corrected to the middle image in series^43^, skull stripped^44^, and coregistered to the T1 structural image. The T1 image was warped to the MNI152 T1 template^45^, and the combined spatial transformation was applied to the mean EPI image. Out of curiosity, the values at each voxel were regressed against the participants’ age using FSL’s^46^ Feat ^47^, and mean EPI values within the thresholded z-stat mask were plotted against age for visual inspection.

### Independent components analysis in larger NKI-Rockland Sample Initiative

As a further exploratory examination, we replicated the preprocessing applied to the fMRI and structural images described in the Preliminary Observation section, above. However, we expanded the sample size to include 1354 cross-sectional participants from the NKI-RS ranging from 6 to 86 years of age (mean = 41.26 +/- 21.43; 836 females).

#### rcFe image creation

All rcFe images were generated with the following procedures. The fMRI sequence was motion corrected^43^, averaged across time, and skull stripped to remove non-brain tissue^44^. The mean skull-stripped image was then z-scaled (zero-meaned and unit variance normalized within the 3D volume mask) to account for any differences in overall image intensity scaling. The images were then sign-inverted so that higher numbers reflected putative increases in relative iron concentration. We termed the resulting voxel map a relative concentration of iron (rcFe) image. This image was smoothed with a 4mm iso FWHM Gaussian kernel and forwarded for further analysis.

#### Study-specific template

Prior to spatial normalization of the rcFe image, we created a study-specific template by warping each participant’s T1 image to the MNI 152 2mm T1 template and subsequently averaging across each participant’s image. The spatial transformation from native EPI space to template space was achieved by applying the by combined spatial transforms from the mean skull-stripped EPI to the high resolution T1 structural MRI ^43^, and the transformation from T1 structural MRI study-specific template^45^.

#### Independent components analysis

The spatially smoothed and normalized rcFe images were submitted to independent components analysis via FSL’s^46^ MELODIC, allowing the algorithm to select the optimal number of components. The resulting components were inspected for correspondence with iron-rich subcortical brain structures.

### Aqueous iron phantom

To examine the potential sensitivity of standard fMRI sequences to biologically plausible iron levels, an aqueous iron phantom was created using 50ml plastic tubes, each filled with distilled water and titrated with commercially available monocrystalline iron oxide nanoparticle solution (AMAG Pharmaceuticals, Inc, Waltham, MA) to achieve solutions of 5, 10, 15 and 20 micromolar iron concentrations. These tubes were submerged in a 250ml beaker of distilled water prior to imaging. 20 fMRI volumes were collected using the standard NKI-RSI 1400ms MB4 sequence. These images were motion corrected^43^ and averaged across time to create a mean volume. This volume was masked to include only the phantom and its contents. The values within the mask were z-transformed to scale the values within the volume, and sign-inverted so that higher values might reflect higher iron concentration. Four regions of interest containing 64 contiguous voxels (8×8×8) were created near the spatial center of each vial, and the values at each voxel were extracted and plotted against the millimolar iron concentration for each vial.

### In-Vivo comparison between rcFe and formal iron methods

We next examined the correspondence between fMRI sensitivity to iron concentration and more standard measures of brain iron. Five volunteers participated in this experiment, aged 28 to 68 yoa, 3 males. Following consenting procedures, they were asked to remain motionless while scanned in several MRI protocols. Siemens Clinical (TR=2500ms; TE=19.0ms, 136ms, 252ms, 64 slices; FOV=210; acquisition voxel size=1.0×0.8×2.0mm; duration=5min), 10-echo R2 (TR=3910ms; TE=13.2ms, 26.4ms, 39.6ms, 52.8ms, 66.0ms, 79.2ms, 92.4ms, 105.6ms, 118.8ms, 132ms, 28 slices; FOV=210; acquisition voxel size=1.1×1.1×3.0mm; duration= 7min), and a 6-echo R2* (TR=2223ms; TE=5.92ms, 25.16ms, 30.32ms, 35.48ms, 40.64ms, 45.80ms, 44 slices; FOV=220; acquisition voxel size=1.1×0.9×3.0mm; duration=4min) imaging protocol as representative iron imaging protocols, as well as the NKI-RSI standard 1400ms fMRI protocol TR=1400ms; TE=30ms, alpha=80, 64 slices; FOV=224; acquisition voxel size = 2.0mm iso, multiband factor = 4; duration = 1min), and T1 (MPRAGE; TR=2500ms; TE=3.5ms; TI=1200ms; voxel size: 1.0mm iso; alpha=8; 174 slices, FOV=256mm; Grappa=2) structural imaging protocol.

#### rcFe processing

rcFe images were calculated, as described above, for each participant in the sample, then smoothed with a 4mm iso gaussian spatial kernel and forwarded to further analysis. The spatial transformation from native EPI space to template space was achieved by applying the by combined spatial transforms from the mean skull-stripped EPI to the high resolution T1 structural MRI ^43^, and the transformation from T1 structural MRI study-specific template^45^.

#### Iron Image Preprocessing

Image series were initially masked for in-brain content by automated skull-stripping of the shortest TE image in each sequence, followed by manual inspection and mask editing. A monoexponential decay function was fit at each voxel against echo time within the brain mask to estimate T2 (10-echo sequence) and T2* (6-echo sequence) decay. R2 and R2* images were created by calculating the reciprocal of the T2 and T2* estimates. The combined transform from the native-space lowest TE to high-resolution T1 structural image ^43^, and the T1 to MNI 152 2mm T1 template transformation ^45^ was subsequently applied to the R2 images.

#### ROI creation

We created 10 subcortical ROIs derived from a-priori masks provided with the Harvard-Oxford subcortical atlas provided with FSL^46^. Atlas probability maps were extracted for the nucleus accumbens, caudate, putamen, globus pallidus, and thalamus for the left and right hemisphere separately, thresholded at a 75% probability level and binarized, yielding 10 distinct subcortical ROIs. These ROI masks were used to extract mean regional rcFe and iron content estimates, which were then forwarded for direct comparison.

#### Examination of rcFe clinical relevance within the ADHD200 Sample

To examine the potential of rcFe images to replicate well-established findings regarding diminished brain iron levels in ADHD participants, as well as associations with ADHD clinical features, we took advantage of the publicly-available ADHD200^48^. Specifically we downloaded the NeuroBureau’s Athena pipeline preprocessed fMRI data^49^ already coregistered to MNI template space. A total of 617 participants ranging from 7 to 21 years of age (mean = 12.34, +/- 3.20; 392 males), were entered into the analysis. rcFe images were created (see rcFe preprocessing). These rcFe images were then upsampled to 2mm iso resolution, spatially smoothed with a 4mm iso Gaussian kernel and subjected to interrogation at several levels. Phenotypic data including age, sex, handedness, diagnostic status, measures of symptom severity, and IQ were downloaded from the ADHD200 site (http://fcon_1000.projects.nitrc.org/indi/adhd200/). For the purposes of all analyses here, we collapsed across ADHD subtypes (hyperactive/impulsive, inattentive, and combined) to form a single binary ADHD diagnosis. This resulted in 226 participants with an ADHD diagnostic label and 392 TDCs.

#### Conceptual replication of NKI-RSI ICA analysis and ROI creation

As an initial step, we subjected the ADHD200 rcFe images to independent components analysis and manually inspected the resulting component maps for correspondence with iron-rich subcortical brain structures. We identified 3 subcortical ICA components; these roughly corresponded to the thalamus, putamen, and globus pallidus components previously identified in the NKI-RSI sample. Regions of interest were created by thresholding and binarizing these component maps.

#### rcFe sensitivity to ADHD status and symptomatology

Using the subcortical ROIs identified above, we extracted the mean rcFe values within each mask for each participant, and forwarded them to a series of interrogations. We first examined differences in overall rcFe intensity in each of ROIs according to ADHD status (ADHD *vs*. TDC) in a series of between-Ss t-test. We then investigated the utility of subcortical ROI rcFe values to predict ADHD status under logistic regression, evaluating the prediction’s receiver-operator characteristics (ROC) and area under the curve (AUC). Finally, we examined whether subcortical rcFe ROI values were predictive of ADHD symptomatology in ADHD and TDC participants.

### rcFe, age, and IQ in the larger NKI-RSI

We extracted rcFe values for each of 3 subcortical ROIs identified via ICA in the NKI-RSI sample from each of the 1354 participants described previously. These, along with age and the full scale IQ score from the NKI-RSI phenotypic battery, were forwarded for analysis via varying coefficient model framework^41^ predicting the relationship between each of the subcortical rcFe values and IQ performance as a function of age. As a secondary illustration of the pattern, we split the sample into three groups by age: child (6-17), adult (18-45), and older (46-86), and computed the correlation between mean rcFe ROI values and IQ.

